# Failure to Replicate a Genetic Signal for Sex Bias in the Steppe Migration into Central Europe

**DOI:** 10.1101/114124

**Authors:** Iosif Lazaridis, David Reich

## Abstract

Goldberg et al.(1) used genome-wide ancient DNA data (2) from central European Bronze Age (BA) populations, and their three ancestral sources of steppe pastoralists (SP), Anatolian farmers (AF), and European hunter-gatherers (HG), to investigate whether the SP migration into central Europe after 5,000 years ago (3, 4) was sex biased. By estimating a lower proportion of SP ancestry on the X-chromosome (36.6%) which is primarily carried in females than on the autosomes (61.8%), they suggested that the migration involved a ratio of 5-14 SP males for every female.

We attempted to replicate this finding using *qpAdm* (3), which leverages allele frequency correlations between the admixed (BA) and source (SP, AF, HG) populations with distant outgroups to eliminate potential biases due to genetic drift between the true source populations and the ones used as surrogates for them. Our outgroups are: Mota (5), Ust_Ishim (6), Kostenki14 (7), GoyetQ116-1 (7), Vestonice16 (7), MA1(8), AfontovaGora3 (7), and Levantine Neolithic farmers (9). We ran *qpAdm* with allsnps: YES and Mota as the basis population. The model fits the data (P=0.072 autosomes, P=0.747 chromosome X). For the BA population, we estimate 61.4±2.9% SP, 31.0± 1.2% AF, 7.6±2.9% HG ancestry using all autosomal SNPs and 67.5±17% SP, 26.5±6.9% AF, 6.0±16.4% HG using all X-chromosome SNPs; thus we do not find less SP ancestry on the X-chromosome.

To diagnose why we do not replicate the finding of (1), we simulated pseudo-BA individuals with known admixture proportions and then used supervised ADMIXTURE (10) - the method used in (1) - on the same 4,605 X-chromosome SNPs. We started our simulations using all possible combinations of (20 AF) x (9 HG) x (8 SP) individuals. For each of these 1,440 triples we simulated 16 individuals, randomly sampling alleles from the source individuals according to the *qpAdm*- estimated proportions of the 16 real BA individuals, and removing simulated individuals with fewer than 1,000 SNPs as in (1) (results were little changed with no filtering or a 2,000 SNP threshold). We estimated ancestry for each individual by performing 10 random-seeded supervised ADMIXTURE replicates and averaging the results (we used the remaining 19 AF, 8 HG, 7 SP individuals as source populations) (Figure 1a). Supervised ADMIXTURE predicts real ancestry poorly in this setting (Figure 1b). The estimation error (estimated - real ancestry) is strongly correlated (*r*=0.91) with the estimated SP ancestry, allowing us to predict it by a regression (Figure 1c), which indicates upward bias for high SP ancestry estimates and downward bias for low ones. For the pool of 16 individuals that the authors of (1) estimated had 36.6% ancestry, the error is predicted to be −19.5%, largely eliminating the discrepancy between the X-chromosome estimate of (1) and the autosomal ancestry estimates of ADMIXTURE (1) and *qpAdm.* These results show that bias in the estimation of admixture proportions, rather than sex bias in the steppe migration, drives the findings of (1).

**Figure 1:**
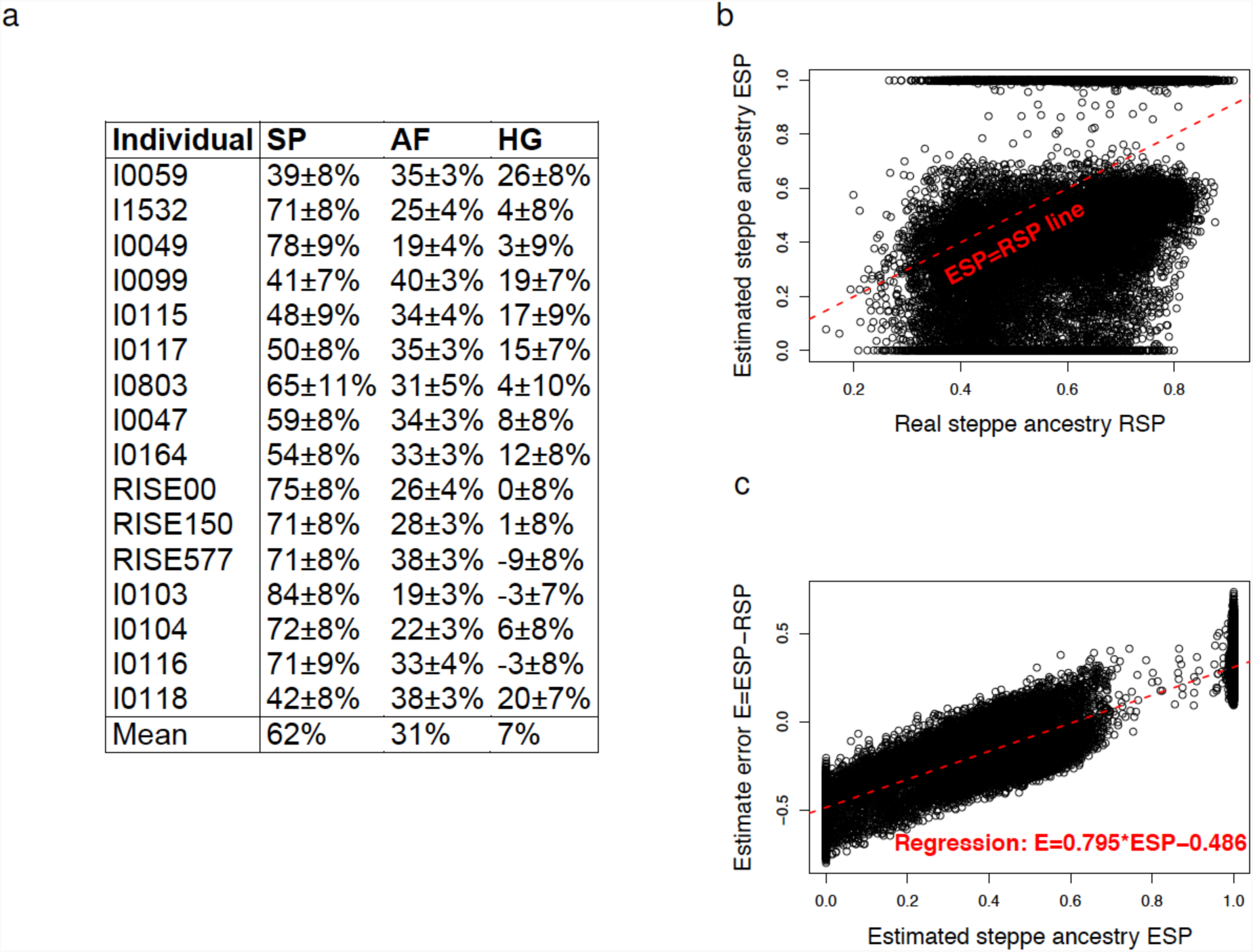
(a) *qpAdm* estimates of autosomal ancestry of 16 BA individuals. (b) Real X-chromosome steppe ancestry (RSP) is not well-predicted by supervised ADMIXTURE-estimated X-chromosome steppe ancestry (ESP) (c) Error (ESP-RSP) is correlated with ESP, allowing us to infer a 19.5% downward bias at the 36.6% ESP ancestry level.

## Acknowledgements

We thank members of the Reich laboratory for critical comments and the authors of (1) for sharing data and program output with us. This work was supported by NIH grant GM100233 and by NSF HOMINID grant BCS-1032255. D.R. is an investigator of the Howard Hughes Medical Institute.

## References

1. Goldberg A, Günther T, Rosenberg NA, & Jakobsson M (2017) Ancient X chromosomes reveal contrasting sex bias in Neolithic and Bronze Age Eurasian migrations. Proceedings of the National Academy of Sciences 114(10):2657–2662.

2. Mathieson I, et al. (2015) Genome-wide patterns of selection in 230 ancient Eurasians. Nature (in press).

3. Haak W, et al. (2015) Massive migration from the steppe was a source for Indo-European languages in Europe. Nature 522:207–211.

4. Allentoft ME, et al. (2015) Population genomics of Bronze Age Eurasia. Nature 522(7555): 167–172.

5. Llorente MG, et al. (2015) Ancient Ethiopian genome reveals extensive Eurasian admixture in Eastern Africa. Science 350(6262):820–822.

6. Fu Q, et al. (2014) Genome sequence of a 45,000-year-old modern human from western Siberia. Nature 514(7523):445–449.

7. Fu Q, et al. (2016) The genetic history of Ice Age Europe. Nature 534(7606):200–205.

8. Raghavan M, et al. (2014) Upper Palaeolithic Siberian genome reveals dual ancestry of Native Americans. Nature 505(7481):87–91.

9. Lazaridis I, et al. (2016) Genomic insights into the origin of farming in the ancient Near East. Nature 536(7617):419–424.

10. Alexander D & Lange K (2011) Enhancements to the ADMIXTURE algorithm for individual ancestry estimation. BMC Bioinformatics 12(1): 246.

